# Auditory processing atypicalities for pure tones and complex speech sounds in Rett Syndrome – towards neuromarkers of disease progression

**DOI:** 10.1101/653246

**Authors:** Olga V. Sysoeva, Sophie Molholm, Aleksandra Djukic, Hans-Peter Frey, John J. Foxe

**Affiliations:** The Cognitive Neurophysiology Laboratory, Ernest J. Del Monte Institute for Neuroscience, Department of Neuroscience, University of Rochester School of Medicine and Dentistry, Rochester, New York, USA; The Cognitive Neurophysiology Laboratory, Departments of Pediatrics and Neuroscience, Albert Einstein College of Medicine & Montefiore Medical Center, Bronx, New York, USA; The Rett Syndrome Center, Department of Neurology, Montefiore Medical Center & Albert Einstein College of Medicine, Bronx, New York, USA; The Laboratory of Human Higher Nervous Activity, Institute of Higher Nervous Activity and Neurophysiology, Russian Academy of Sciences, Moscow, Russia

**Keywords:** EEG, Auditory Evoked Potential, Neurodevelopmental Disorder, Rett Syndrome, Rare Disease, AEP, Biomarker, Endophenotype, MECP2, Females

## Abstract

Due to severe motor impairments and the lack of expressive language abilities seen in most patients with Rett Syndrome (RTT), it has proven extremely difficult to obtain accurate measures of auditory processing capabilities in this population. Here, we examined early auditory cortical processing of pure tones and more complex phonemes females with confirmed mutation of the MECP2 gene. We recorded high-density auditory evoked potentials (AEP), which allow for objective evaluation of the timing and severity of processing deficits along the auditory processing hierarchy. We compared AEPs of 12 females with RTT to those of 21 typically developing (TD) peers aged 4-21 years, interrogating the first four major components of the AEP (P1: 60-90ms; N1: 100-130ms; P2: 135-165ms; N2: 245-275ms). Atypicalities were evident in RTT at the initial stage of processing. Whereas the initial P1 showed increased amplitude to phonemic inputs relative to tones in TD participants, this modulation by stimulus complexity was absent in RTT. Interestingly, the subsequent N1 did not differ between groups, whereas the following P2 was hugely diminished in RTT, regardless of stimulus complexity. The N2 was similarly smaller in RTT, and did not differ as a function of stimulus type. The P2 effect was remarkably robust in differentiating between groups with near perfect separation between the two groups despite the wide age range of our samples. Given this robustness, along with the observation that P2 amplitude was significantly associated with RTT symptom severity, the P2 has the potential to serve as a biomarker of treatment efficacy.

**Significance statement:** Our study points to dramatic reduction of the P2 component of the auditory evoked potential (AEP) as a potentially reliable biomarker of Rett Syndrome severity, with prospective applicability as an objective readout (neuromarker) of change in functional brain activity following therapeutic interventions administered in the context of clinical trials. Compellingly, the reduction of P2 amplitude in patients with RTT mimics findings in animal models of RTT, providing a translational bridge between pre-clinical and human research.

## INTRODUCTION

Rett Syndrome (RTT) is a neurodevelopmental disorder affecting approximately one in 10,000 live female births, irrespective of place of origin or ethnicity (Hagberg et al., 1983; Laurvick et al., 2006; Wong and Li, 2007). It results from spontaneous mutation in the *MECP2* gene located on the X-chromosome (Amir et al., 1999), and its primary clinical features include severe motor deficits and cognitive impairments. Verbal ability is typically also very restricted or absent, and combined with the motor impairments, this creates major challenges for clinicians and caregivers in evaluating the degree to which individuals with RTT can understand and differentiate the speech signal. Direct recordings of the auditory evoked potential (AEP) using high-density electroencephalography (EEG) provide an easily implemented means by which neuroscientists can directly study the neurophysiological bases of auditory signal processing in RTT, allowing for objective assessment of the severity of potential auditory processing deficits. Clinically, these AEP measures may also prove useful as biomarkers of disease severity and progression, as well as targets against which to measure the efficacy of therapeutic interventions during clinical trials.

Early electrophysiological studies in RTT mostly focused on the auditory brainstem response (ABR), with findings indicating that initial subcortical stages of auditory signal processing appeared mostly unaffected (Stach et al., 1994). In contrast, AEP studies assessing later cortical stages of auditory processing have tended to show quite significant impairments (Bader et al., 1989; Stach et al., 1994; Stauder et al., 2006; Foxe et al., 2016; Peters et al., 2017; Brima et al., 2019; Key et al., 2019). For example, Stach and colleagues (Stach et al., 1994) reported that more than 50% of individuals with RTT showed atypical AEPs, although they did not precisely specify what these abnormalities were. Others have pointed to delayed auditory processing. For example, Bader and colleagues (Bader et al., 1989) reported significant prolongation of the peak latencies of a series of AEP components (i.e. Pa, N1 and P2), and Stauder and colleagues (Stauder et al., 2006) uncovered atypical developmental changes in the latency of the N2. Recent work from our group accords well this notion of slowed auditory processing, as we found substantial delay in the mismatch negativity (MMN) response, an AEP component that is automatically generated when occasional deviant stimuli interrupt a stream of standard stimuli (Foxe et al., 2016). In that study, we specifically assessed MMN responses to large changes in pitch, and the implications of our results were that cortical representation of even such a basic feature as frequency representation was atypical and delayed in RTT. Similarly, using the MMN to assess auditory sensory memory for duration, we found that individuals with RTT showed substantial impairment, such that the duration-evoked MMN was only present at rapid stimulation rates (∼2Hz) but was not detectable when slower stimulation rates (circa 0.5 and 1 Hz) were utilized (Brima et al., 2019).

While there is some AEP research using frequency-specific tone-pip stimuli in RTT syndrome, AEPs in response to more complex sounds such as speech, have not yet been closely examined. Given that a key question in the RTT population is the extent to which non- or minimally- verbal individuals can process and understand the spoken word, there is an imperative to map cortical auditory processing abilities in these individuals. Recent reports by Key and colleagues suggest that there may be quite profound processing atypicalities for speech tokens. They reported abnormalities in the AEP to speech stimuli in the latency range from 200-500 ms post-stimulus when the responses to real words were compared to those to non-words (Key et al., 2019) and from 250-450 ms and 450-750 ms when AEPs to a participant’s own name were compared to those evoked by other names (Peters et al., 2017). However, the work was not designed to assess the earlier stages of cortical processing and the initial component structure of the AEP as relatively few trial repetitions were recorded (n = 32), which precludes reliable differentiation of earlier AEP components. As such, there remains a need to systematically study the initial sequence of AEP components (i.e. P1, N1, P2 and N2) in response to speech tokens, so that the integrity of the successive processing stages along the auditory hierarchy can be examined.

Here, we use high-density electrical mapping of the AEP to probe these processing stages in response to both simple tone-pip and complex phonemic sounds in females with RTT. To ensure high signal-to-noise ratios, and thus allow for detailed interrogation of the response, sound stimuli in our experimental were repeated more than 800 times each.

## METHODS

### Participants

Thirteen female patients with Rett syndrome participated in this study (mean age 12.9; range 3.9–20.6). They were recruited during clinical visits to the Rett Center at the Children’s Hospital of Montefiore Medical Center in the Bronx, New York. Diagnosis was based on current diagnostic criteria (Neul et al., 2010) and was confirmed clinically by a medical doctor specializing in this population (author A.D.) as well as via genetic testing. Symptom severity was assessed for each patient using the Rett Syndrome Severity Scale (RSSS), as modified by Kaufmann and colleagues (Kaufmann et al., 2012). This clinician-rated scale represents an aggregate measure of the severity of clinical symptoms, including motor function, seizures, respiratory irregularities, ambulation, scoliosis, and speech. Each item is scored from 0 (absent/normal) to 3 (severe). Demographic characteristics of the Rett group are summarized in Table 1. RTT participants were compared to a control group of 21 age-matched females (mean age 12.44; range 4.3–21.1) who were all typically developing, with no familial history of Rett syndrome and no current or lifetime history of psychiatric, neurodevelopmental, or neurological disorders. The study groups were of similar age (t (31) = 0.32, p = 0.75). This study was approved by the institutional review board of The Albert Einstein College of Medicine (Protocol Reference Number #2011-447). Written informed consent was obtained from parents or legal guardians, and where possible, assent from the patient was also ascertained. All aspects of the research conformed to the tenets of the Declaration of Helsinki.

**Table 1.** RTT cohort characteristics. Each participant’s age (years), scores on the Rett Syndrome Severity Scale (RSSS), comorbid seizure diagnosis and ability to walk (ambulation) are listed. Note that RTT participant #3 was excluded from the main analysis due to high amplitude noise in the EEG that precluded derivation of an adequate AEP (indicated by the asterisk).

| RTT | Age | RSSS | Seizures | Ambulation |
| --- | --- | --- | --- | --- |
| 1 | 3.9 | 7 | No | No |
| 2 | 5.1 | 13 | No | Yes |
| 3 | 6.9* | 14 | No | No |
| 4 | 8.6 | 14 | Yes | Yes |
| 5 | 10.1 | 12 | No | Yes |
| 6 | 13.0 | 15 | Yes | No |
| 7 | 13.5 | 14 | Yes | No |
| 8 | 13.8 | 16 | Yes | No |
| 9 | 13.9 | 10 | Yes | Yes |
| 10 | 16.3 | 19 | Yes | No |
| 11 | 16.9 | 12 | Yes | Yes |
| 12 | 20.1 | 6 | No | Yes |
| 13 | 20.6 | 13 | Yes | No |

### Experimental design

Participants sat in a darkened sound-attenuated electrically shielded booth (Industrial Acoustics Company, Bronx, NY), either in a chair/wheelchair or on a parent’s lap, while watching a movie of their choice on a laptop (Dell Latitude E640) with the volume turned off. Auditory stimuli were presented using a pair of speakers (Bose Companion 2 Series II, Multimedia Speaker System) placed behind the laptop. An oddball paradigm was employed whereby standard and deviant auditory stimuli were presented randomly with a probability of 0.85 to 0.15, respectively. There were two experimental conditions with simple tones and phonemes, respectively. In the “Tone” condition, stimuli were sinusoidal tones of two different frequencies, 1000 Hz for the standards and 500 Hz for the deviants. Both tones had a duration of 100 ms, a rise and fall time of 10 ms, and an intensity of 75 dB SPL. In the “Phoneme” condition, the phoneme /ba/ served as the standard and the phoneme /da/ served as the deviant. Phoneme duration was 250 ms, and they were presented at an intensity of 65 dB SPL. Here we report the data only in response to the more frequently presented standard stimuli, comparing them across stimulus conditions. The time-frequency characteristics of the stimuli are shown in Figure 1. The Rett group completed an average of 9.64 and 9.53 blocks (range 7– 11), with each block containing 140 stimuli, which were presented with a stimulus-onset-asynchrony (SOA) of 900 ms, while the control group completed an average of 10.04 and 9.95 blocks (range 10–11), for Tone and Phoneme conditions respectively. Simple tone blocks were mostly presented first, except in the case of one RTT participant and 8 TD cases. Block order did not influence the AEPs of interest in the TD sample as examined by ANOVA: Type by Condition Order interaction (F(1,19)< 2.47; p >0.13 for all components).

**Figure 1:**
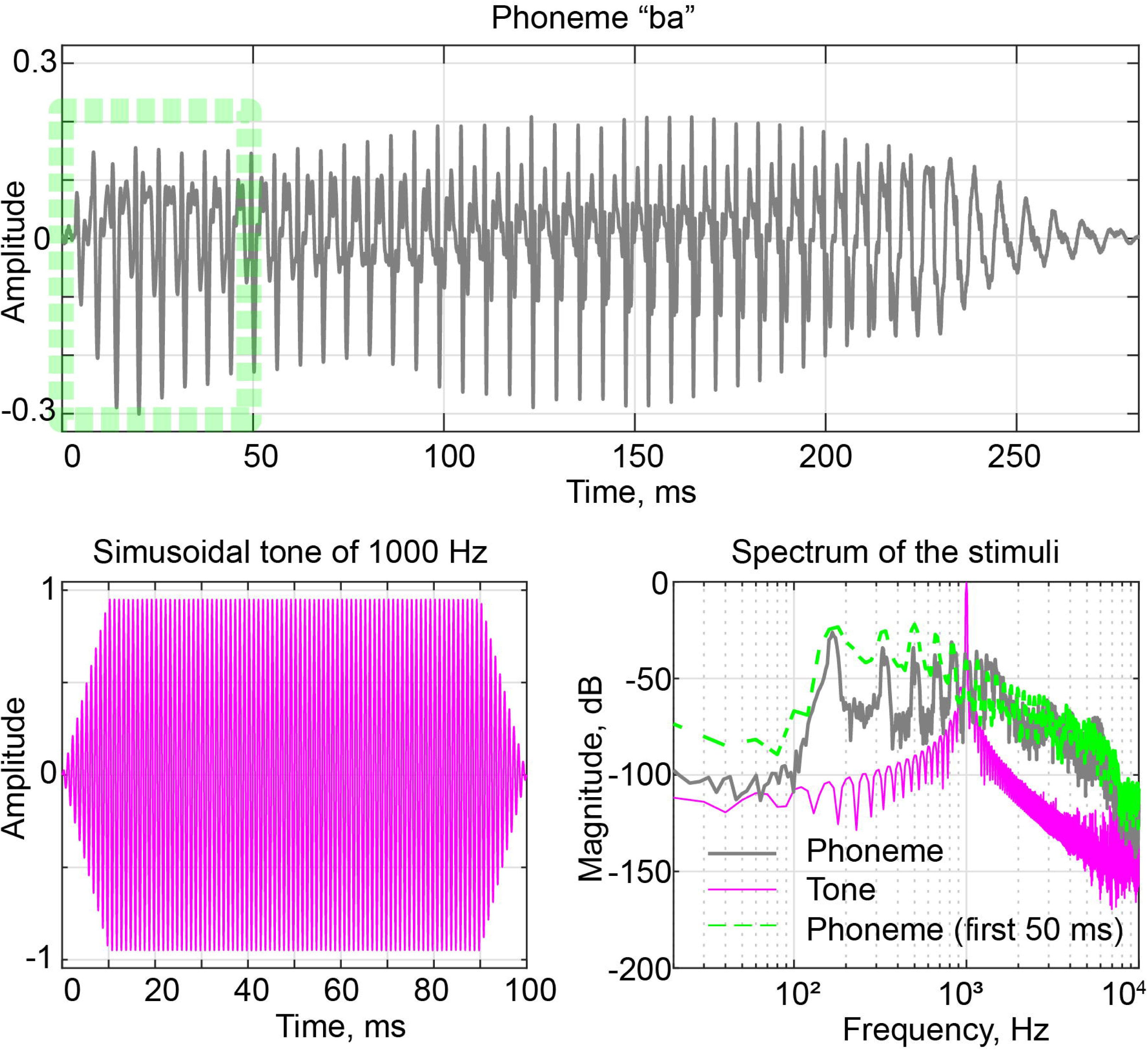
Experimental stimuli. Top: Waveform of phoneme /ba/. Lower Left: Waveform of sinusoidal tone of 1000 Hz. Lower right: Spectrum of the stimuli (blue for Phoneme, red for Tone and green for the first 50 ms of Phoneme).

### EEG recordings

Continuous EEG data were recorded using a Biosemi ActiveTwo 64 electrode array, analog-to-digital converter, and fiber-optic pass-through to a dedicated acquisition computer (digitized at 512 Hz; DC-to-150 Hz pass-band). Biosemi replaces the ground electrodes that are used in conventional systems with two separate electrodes: common mode sense and driven right leg passive electrode. These two electrodes create a feedback loop, thus rendering them as references.

### Data processing

All EEG processing and analyses were performed in MATLAB (the MathWorks, Natick, MA, USA) using custom scripts and the FieldTrip Toolbox (Oostenveld et al., 2011). Following recording, the continuous EEG was segmented into epochs of 800 ms in length, from −200 to +600 ms post-stimulus latencies. All epochs were arranged by stimulus type and concatenated. Than the data were bandpass filtered 1-20 Hz using zero-phase forward and reverse Butterworth infinite impulse response filter and examined for artifacts. Artifact identification was done via z-scores and variability measures: all trials that deviated more than 2 standard deviations from average were removed. Noisy and bad channels were also identified based on the same algorithm and then restored by interpolation from neighboring channels. The AEP of one RTT participant contained an unacceptable level of high amplitude noise that was more than ten times that which was observed in the typical AEP amplitude, and this participant was therefore excluded from further analysis. After artifact rejection, the mean numbers of trials for the RTT and TD groups respectively in the Tone condition were 756 and 804, and in the Phoneme condition 733 and 763 (ranging from 558 to 906). AEPs were calculated separately for each condition by averaging all relevant trails. Epochs were baselined to the 200 ms pre-stimulus interval and re-referenced to the average of TP7 and TP8 sites (which fall over the left and right mastoid regions where the auditory ERPs tends to invert in polarity vis-à-vis the fronto-centrally focused response). The main contributions to the major AEP components (P1, N1, P2 and N2) come from generators in and around primary auditory cortex along the supra-temporal plane and as such, they give rise to predominantly fronto-central scalp distributions. Thus, for peak amplitude analyses here, we focused on the averaged AEPs from the FCz, FC3 and FC4 electrodes.

### Statistical analysis

To statistically characterize potential between-group differences, we used SPSS software (IBM SPSS Statistics version 25). Averaged amplitudes of the AEP components within the following characteristic latency windows indentified based on TD grand-averaged AEPs (P1: 60-90 ms, N1: 100-130 ms, P2: 135-165 ms, and N2: 245-275) were entered into an analysis of variance (ANOVA) with group (RTT vs TD) as a between-subjects factor and stimulus-type (Tone vs Phoneme) as a within-subjects factor.

It is important to note that this analysis is not sensitive to potential differences in the latencies of these AEP components, a factor we were interested in given the prior evidence for slowing of AEP responses. Therefore, we conducted supplementary analyses by identifying the AEP components for each individual in the Tone condition, where the early AEP components are more pronounced and dissociable. For each individual, the P1, N1, P2 and N2 components of the AEP were identified via the following automatic procedure. P1 was estimated as the first positive component elicited at least 40 ms after onset time, N1 as the first negative component detected in the 80-200 ms latency range. The second positive component was labeled P2, and N2 was defined as the negative peak with the greatest negative-going value in the 200-400 ms range. This automatic detection algorithm corresponds well with manual peak detection. One RTT subject had no clearly detectable peaks and was excluded from further analysis. In addition, one 21-year old TD subject had no detectable P1 component and three subjects (two TD participants aged 4.3 and 6.9 years, as well as one RTT aged 10.10 years old) had no evident N1 component. Technically, P2 could be defined in these subjects without the N1 component, but the P2 latencies deviated significantly from the rest of the sample, at least when the TD group was considered. Thus we also excluded them from the final analysis. Student t-tests were used to examine the between-group differences in the latencies and amplitudes of AEPs in this supplementary analysis.

Pearson correlations were used to assess potential relationships between AEP components and age. We also compared the correlation coefficients for each component between groups using a Fisher z-transformation to examine if AEP maturation was similar across groups (Meng et al., 1992). The AEP amplitudes of interest that showed between-group differences were also assessed for potential correlation with Rett Syndrome Severity Scores in the participants with RTT.

## RESULTS

The grand-averaged AEPs in response to both tones and phonemes showed the expected pattern of identifiable P1, N1, P2 and N2 components in the TD participants (Figure 2). AEPs in the RTT group, however, clearly deviated from those of TDs, with direct correspondence of component peaks being difficult to discern for all but the initial P1.

**Figure 2.**
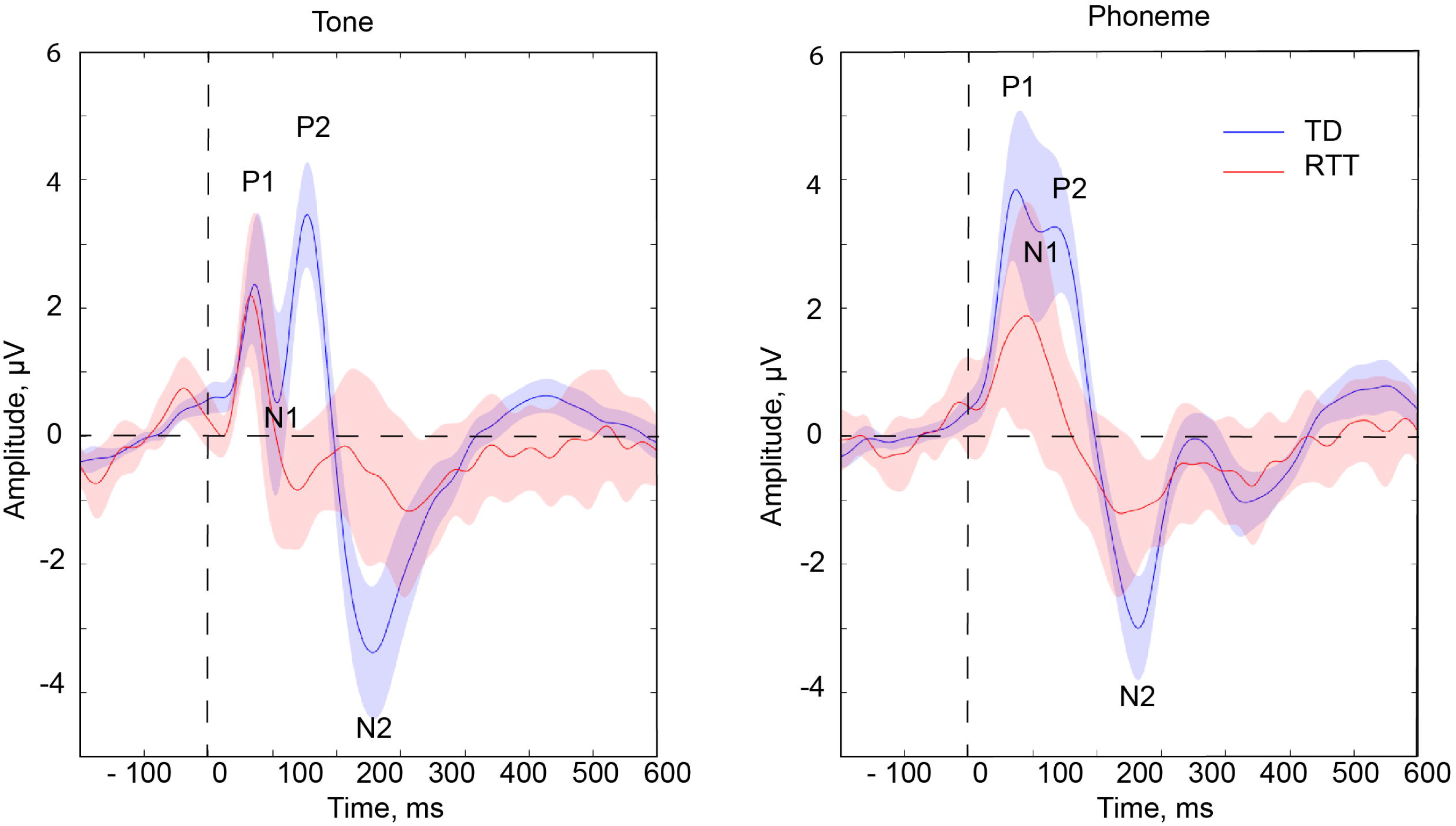
Auditory Evoked Potentials (AEPs) in response to Tones and Phonemes in TD and RTT groups. The vertical dashed line corresponds to the onset of the stimuli, and the horizontal dashed line indicates baseline value. Blue and red lines correspond to TD and RTT AEPs respectively. Opaque blue and red shading illustrates the standard error of the mean.

Results of the ANOVA are summarized in Table 2. A significant Stimulus-type by Group interaction was observed for the P1 component, and this stemmed from the significantly greater amplitude of the P1 in response to phonemes as compared to tones (t(20) = 5.185; p < 0.0001) in the TD group, an effect that was not present in the RTT group (t(11) = 0.046; p = 0.964, Figure 3). The N1 amplitude was reduced in response to phonemes compared to tone stimuli, similarly across groups, with a significant main effect of stimulus-type, but no significant Group effect nor interaction of Stimulus-type by Group.

**Table 2.** ANOVA effects. The numbering convention lists the F-Statistic value, followed by the p-value, followed by the effect size expressed as eta-squared (i.e. F/p/η2). An asterisk is used to further denote cells with significant findings.

|  | Stimulus Type | Group | Group X Stimulus-Type |
| --- | --- | --- | --- |
| P1 60-90 ms | 8.996/0.005/0.225* | 1.503/0.229/0.046 | 8.518/0.006/0.216* |
| N1 100-130 ms | 43.026/<0.0001/0.581* | 2.812/0.104/0.083 | 0.508/0.481/0.016 |
| P2 135-165 ms | 0.945/0.338/0.030 | 36.928/<0.0001/0.544* | 2.681/0.112/0.080 |
| N2 245-275 | 0.019/0.892/0.001 | 10.756/0.003/0.258* | 1.614/0.213/0.049 |

**Figure 3.**
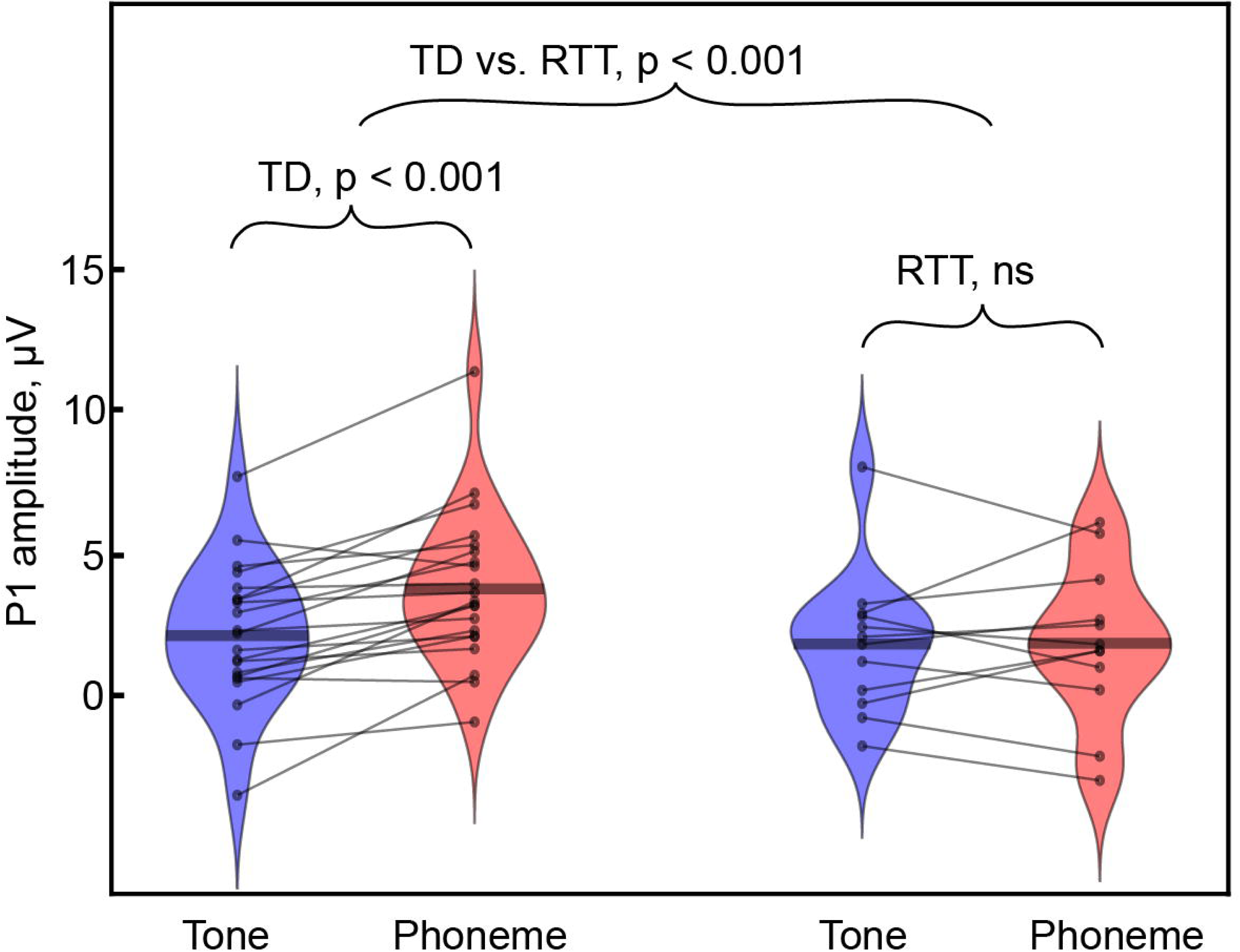
Modulation of P1 amplitude by stimulus type in TD (left) and RTT (right) groups. Each individual’s data are represented by connected dots corresponding to values in the Tone and Phoneme conditions. Note the significantly smaller P1 in the Tone than in the Phoneme condition in TD, a pattern that is not observed in the RTT group.

In contrast to the PI and N1, the P2 component differed substantially between the RTT and TD groups, but was not modulated by stimulus type (Table 2). As can be seen in Figure 2, whereas there is a clear and prominent P2 component in the TD group for both Tones and Phonemes, at the group level it appears to be entirely absent in participants with RTT. The lack of statistical difference between P2 in response to Phonemes and Tones suggests that the P2 amplitudes in these two conditions should be considered together, thus we averaged P2 amplitude across conditions for further exploration. Consideration of individual subject data (Figure 4) reveals that only three RTT patients had P2 amplitudes exceeding the minimum value observed for TD participants (and these were still smaller than the average across TD participants). A significant Pearson correlation between RSSS and P2 amplitude suggests relevance of the P2 component to clinical RTT manifestation (r(12) = −0.62, p = 0.032, Figure 5). By placing a decision threshold at a P2 amplitude of + 0.63 *µ*V, RTT patients can be distinguished from controls with 100% specificity and 75% sensitivity.

**Figure 4.**
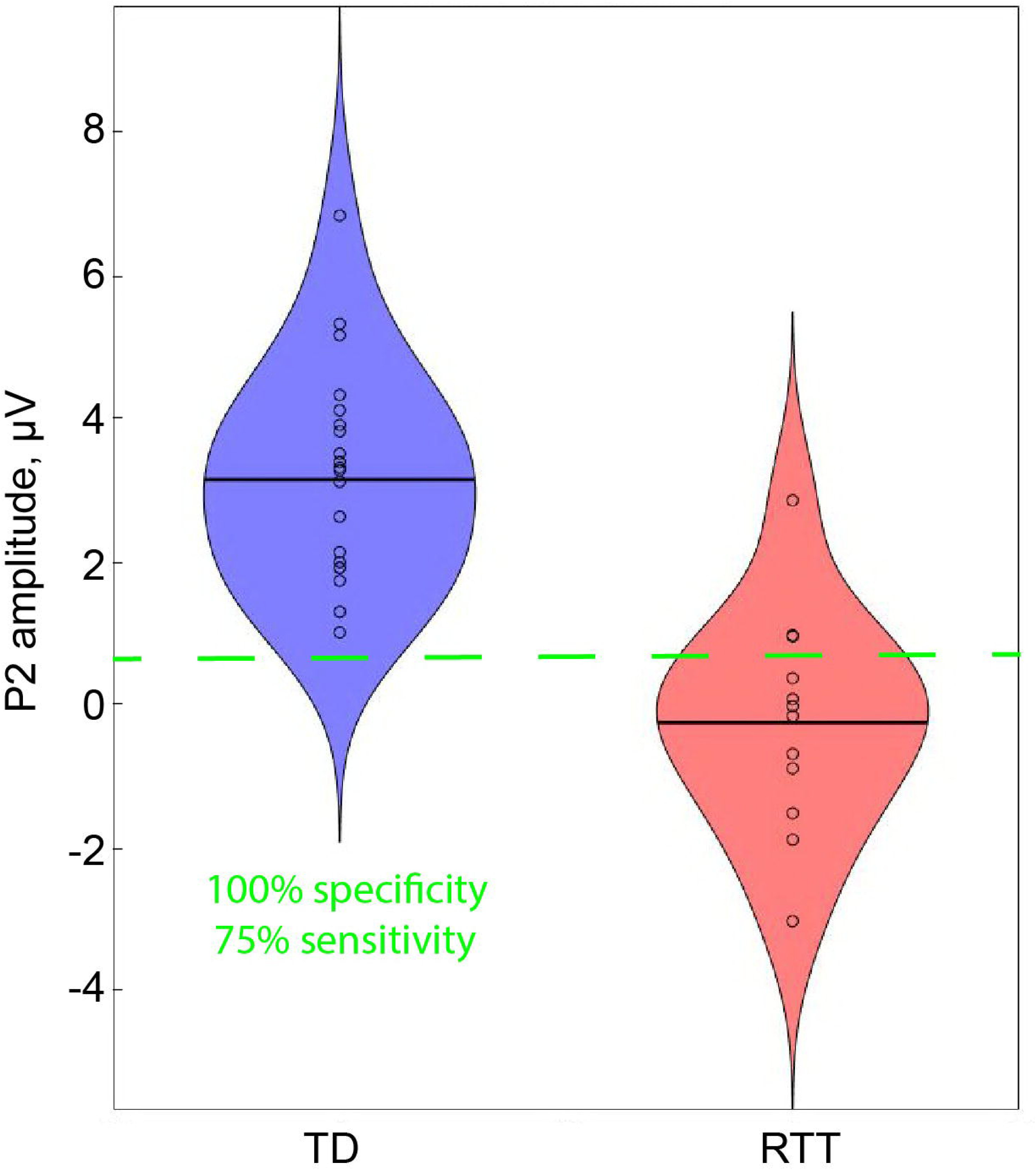
Amplitude of the P2 component in TD and RTT groups. The figure represents data averaged across conditions (Tone and Phoneme). Each dot corresponds to an individual participant. Two short horizontal lines indicate the means of the group values. The dashed horizontal green line is drawn at the decision threshold of + 0.63 µV, where RTT and TD participants are discriminated with 100% specificity and 75% sensitivity.

**Figure 5.**
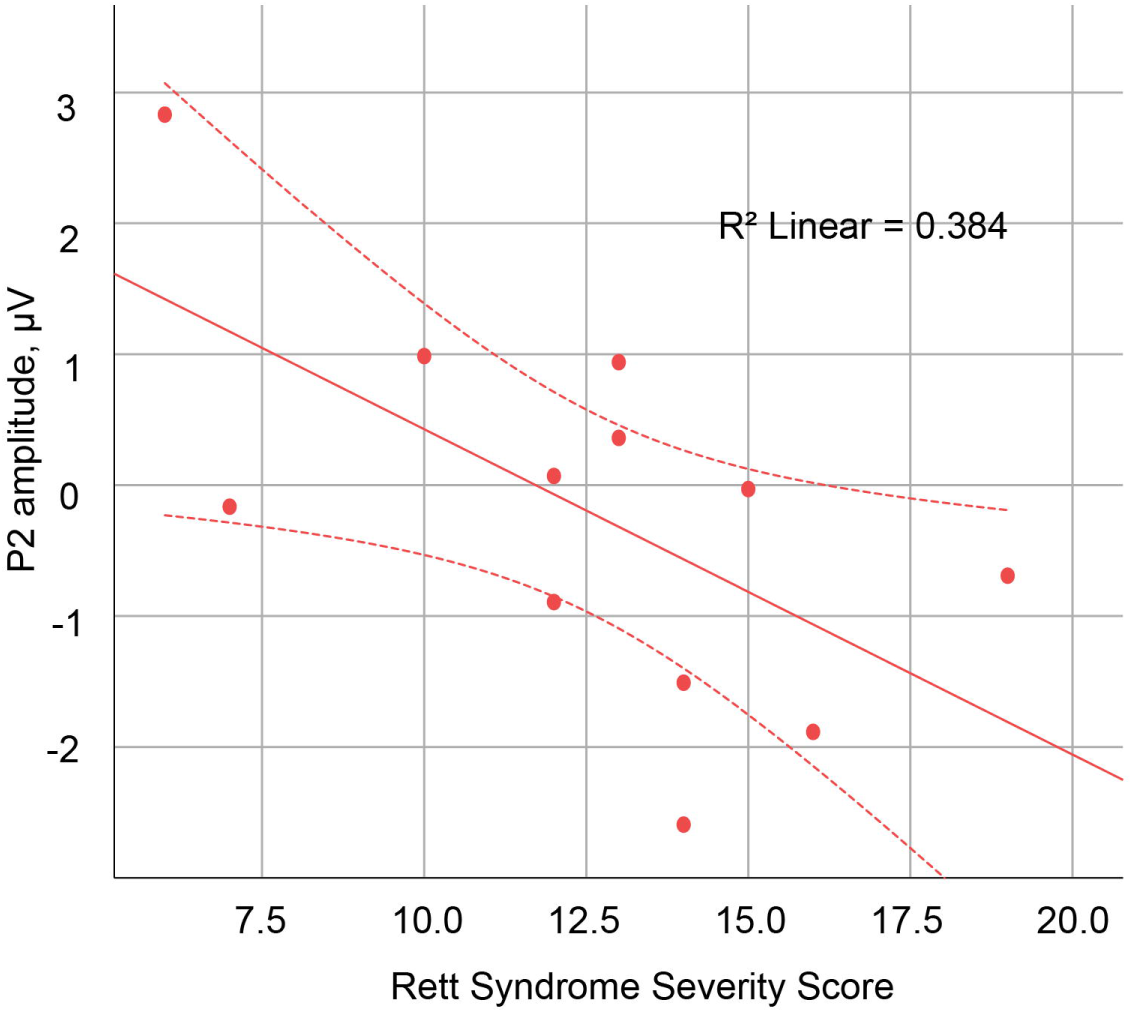
Relationship between P2 component and Rett Syndrome Severity Scores (RSSS) in patients with RTT. Each dot represents data from an individual participant. Note that larger RSSS values indicate more severe symptoms.

The N2 response was also significantly reduced in RTT (main effect of group, Table 2, Figure 6). However, the result is a less robust indicator of group membership than the P2 component: only about 50% of RTT patients’ N2 amplitudes lie below the minimum N2 amplitude value seen in the TD group.

**Figure 6.**
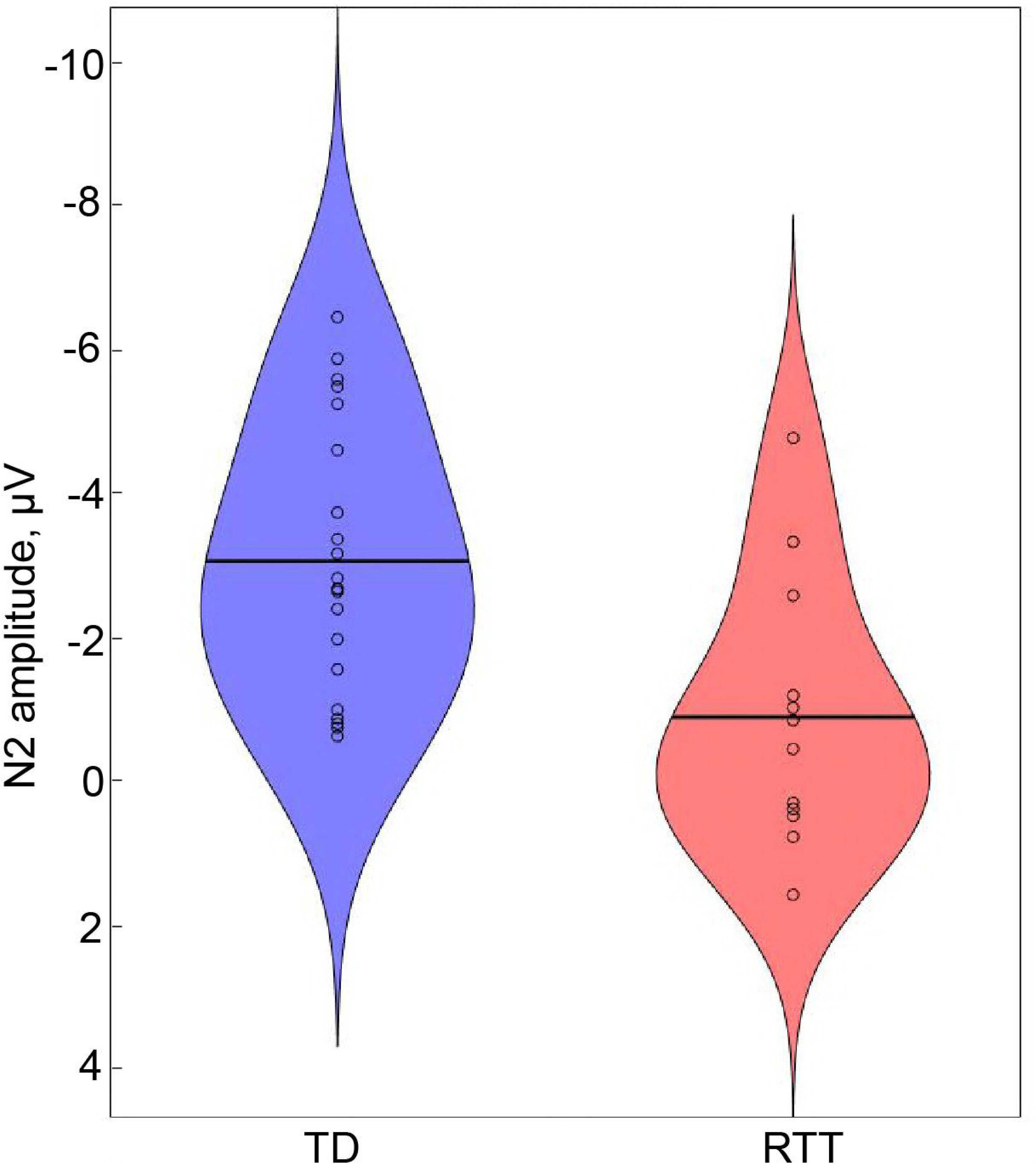
Amplitude of N2 component in TD and RTT participants. The figure represents data averaged across conditions (Tone and Phoneme). Each dot corresponds to an individual participant. Two short horizontal lines indicate the mean of each group.

**Figure 7.**
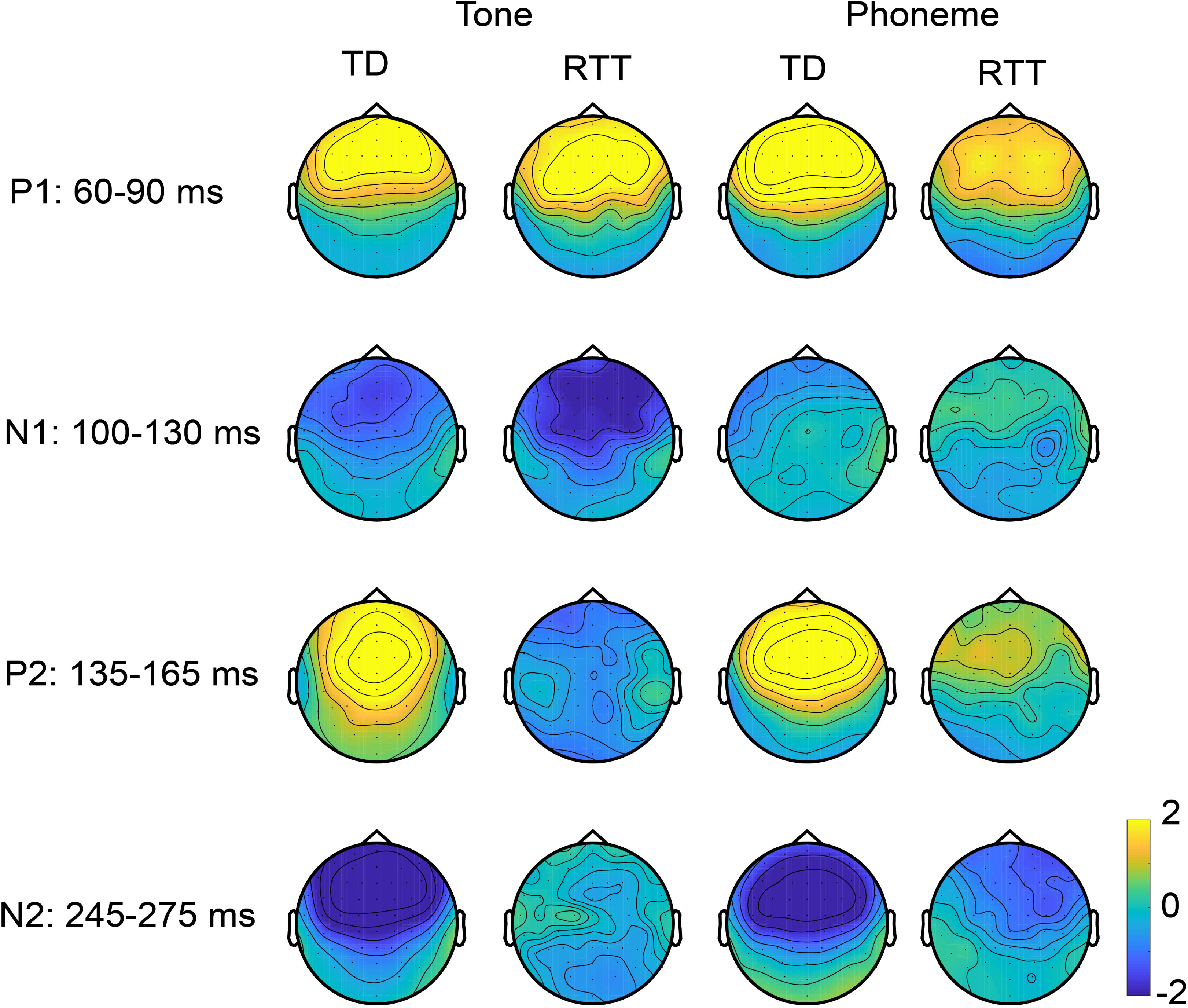
Topography of auditory evoked potentials in response to Tone and Phoneme conditions in TD and RTT groups. While topography for P1, P2 and N2 components are plotted based on averaged amplitude over corresponding time intervals, N1 topography is plotted in relation to P1 meaning that averaged over N1 latency AEP amplitudes were subtracted from AEP amplitudes averaged over P1 latency interval.

Topographic voltage maps suggest a similar fronto-central scalp distribution for the P1 component irrespective of stimulus type and subject group. The N1 topography when plotted relative to P1 shows a clear fronto-central negative focus for the Tone condition in both groups, while in Phoneme condition it was not particularly evident due to low response strength. The P2 and N2 topographies, in contrast, showed clearly distinguishable spatial distributions between groups, with a fronto-central maximum observed only in the TD group, while maps for the RTT group were clearly atypical (Figure 6).

### Analysis of AEP latencies in the Tone condition

The results of this analysis confirmed the results of the primary analysis. No significant between-group differences were evident for P1 and N1 components, with their amplitudes and latencies essentially being typical in patients with RTT (Table 3). P2 and N2 components were significantly attenuated as well as delayed in patients with RTT. However, P2 latency delay was much less pronounced than the P2 amplitude reduction outlined above.

**Table 3.** Results of RTT vs. TD group comparison in Tone’s AEPs characteristics. Number indicate the significance level (p-value) of Student t-test comparison.

|  | Latency | Amplitude |
| --- | --- | --- |
| P1 60-90 ms | 0.227 | 0.682 |
| N1 100-130 ms | 0.065 | 0.320 |
| P2 135-165 ms | 0.004* | 0.00005* |
| N2 245-275 ms | 0.000004* | 0.0212* |

### Developmental changes in the AEP

Our study confirmed developmental decreases in P1 and N2 amplitudes and increases in N1 strength in the TD group but was not adequately powered to detect between-group differences in the developmental trajectory of AEP components (Table 4).

**Table 4.** AEP developmental changes. Numbers in the second and third columns are the Pearson correlation coefficients (r) and their significance values (p) for correlations between age and the amplitudes of the four AEP components of interest. Fisher’s z transformation and its significance value (p) were used to compare the correlation coefficients between the groups and are represented in the fourth column. An asterisk is used to denote correlations that reached significance. Note that while developmental changes were significant only in the TD group, there were no detectable differences in the correlation coefficients between RTT and TD groups.

|  | TD (n = 21) | RTT (n = 12) | TD vs RTT difference |
| --- | --- | --- | --- |
| P1 60-90 ms | -.631/.002* | -.447/.145 | z = -0.64, p = 0.52 |
| N1 100-130 ms | -.636/.002* | -.183/.569 | z = -1.39, p = 0.16 |
| P2 135-165 ms | -.398/.074 | .184/.566 | z = -1.49, p = 0.14 |
| N2 245-275 | .700/<.0001* | .227/.479 | z = 1.56, p = 0.12 |

## DISCUSSION

This is the first study to examine early auditory cortical processing of complex speech stimuli in RTT, and to compare these responses to those evoked by simple frequency-specific tones. The initial AEP component (the P1 – 60-90 ms), which localizes in the region of primary auditory cortex (Leavitt et al., 2007; Gomez-Ramirez et al., 2011), showed increased amplitude to the more complex phonemic inputs relative to pure tones in TD participants. In contrast, while a relatively robust P1 component was evident in the RTT population, it did not show this modulation by stimulus type/complexity. Interestingly, this effect did not simply cascade up the auditory system. Rather, the subsequent N1 (100-130 ms) did not differ between groups, whereas the P2 (135-164 ms), the following major deflection, was hugely diminished in RTT, regardless of stimulus type. The N2 (245-275 ms) was also substantially reduced in amplitude in RTT, and showed no modulation as a function of stimulus type. The P2 effect was remarkably robust in differentiating between groups with almost perfect classification into group despite the wide age-range of our samples. Given this robustness, the P2 has potential to serve as a biomarker of treatment efficacy. The implications of these findings are discussed in what follows.

The only prior study to compare early ERP components between TD and RTT groups did not observe P2 and N2 reductions in RTT, but rather, reported delayed latencies of the N1 and P2 components in response to simple tones (Bader et al., 1989). The current data did not replicate an N1 latency delay, but confirmed a sizeable P2 delay. These discrepancies between results are likely linked to differential sample characteristics, such as sample size (7 vs 12), age range (10-22, vs 4-22) and possible differences in the genetics of the RTT participants (unknown vs confirmed *MECP2* mutation) in Bader”s and our studies respectively. Another contributing factor may have been lower signal-to-noise ratios in the Bader study due to considerably lower numbers of trials (n=128).

### Neurophysiological differentiation of simple versus complex sounds

#### The P1 component (60-90 ms)

The P1 component, which is generated in early auditory cortex (Liégeois-Chauvel et al., 1994; Leavitt et al., 2007; Ruhnau et al., 2011), is known to be affected by stimulus parameters such as complexity or “speechness” in TD children (Čeponiené et al., 2001; Čeponienė et al., 2005; Čeponienė et al., 2008; Bruder et al., 2011; Kuuluvainen et al., 2016). In particular, P1 amplitude is larger in response to vowels than to simple sinusoidal tones in school-aged children (Čeponiené et al., 2001; Bruder et al., 2011). Čeponiené and colleagues suggested that this increase in amplitude was related to higher neuronal synchronization in response to phonemes as compared to simple tones (Tervaniemi et al., 2000; Čeponiené et al., 2001), whereas Bruder and colleagues forwarded a somewhat different explanation, positing increased P1 amplitude as a reflection of a wider neural tuning curve (i.e. larger neuronal population activation) to the phonologically richer speech sounds (Bruder et al., 2011). Here, we confirmed P1 amplification in TD children in response to spectrally richer speech sounds compared to tones, a modulation notably absent in the RTT cohort. This lack of modulation in RTT may reflect a deficit in increasing synchronization of neural response to more complex stimuli, or a breakdown in the fundamental tonotopic representation of spectral information within core auditory cortical regions. Our previous finding that the MMN to frequency deviations was both attenuated and delayed in RTT accords well with the latter interpretation, suggesting that even fundamental tonotopic representation in early auditory core regions may be weakened (Foxe et al., 2016).

#### The N1 component (100-130 ms)

The subsequent N1 component is also thought to have its major generators along the supra-temporal plane, in and around primary auditory cortex (Liégeois-Chauvel et al., 1994; Lütkenhöner and Steinsträter, 1998; Leavitt et al., 2007; Ruhnau et al., 2011). In contrast to P1, N1 was larger for tones than for phonemes in both groups. However, despite apparent differences in the AEP during the N1 timeframe of 100-130 ms post-stimulus in the same\adjunct regions, our analysis did not reveal a statistically significant group difference. This may reflect greater variability at the individual participant level at this later stage of auditory processing.

### Robust RTT vs TD differentiation

#### P2 attenuation (135-165 ms)

The P2 component has been localized to several sources in auditory cortex as well as in the non-specific reticular-thalamo-cortical activating system (Knight et al., 1988; Velasco et al., 1989; Lütkenhöner and Steinsträter, 1998; Godey et al., 2001; Crowley and Colrain, 2004). A growing body of evidence has related the P2 component to consolidation processes associated with auditory memory formation and learned relevance (Tremblay et al., 2010; Tremblay et al., 2014). Supporting this link, multiple studies have reported increases in P2 amplitude following perceptual learning/training (Tremblay et al., 2001; Atienza, 2002; Reinke et al., 2003; Bosnyak, 2004), or in populations with increased exposure to particular stimuli, such as musicians (Shahin et al., 2005; Kuriki, 2006; Park et al., 2018). It is also of note that the P2 can show increased amplitude several days after initial exposure to stimuli, even when no training or additional exposure to the stimuli takes place (Sheehan et al., 2005; Tremblay et al., 2014). This P2 augmentation to background familiar stimuli suggests that it reflects automatic integration of previous stimulus history, and does not necessitate an explicit training process *per se*. It is particularly noteworthy that a P2 increase following training or stimulus exposure, does not appear to occur immediately, but rather, is seen only on the next day, and this augmentation can then last for months (Atienza, 2002; Ross and Tremblay, 2009; Tremblay et al., 2014). This pattern of results certainly seems to suggest that the P2 is related to memory consolidation processes and that sleep plays a key role in its production. Additionally, P2 amplitude recorded during passive conditions shows an inverse relationship to behavioral improvements after training, quantified as reaction time speeding in response to the trained stimuli (Tong et al., 2009; Talebi et al., 2015). Such an association between response speed and AEP amplitude points to the relevance of P2 enhancement for effective task performance.

In the current study, the P2 component was drastically attenuated in patients with RTT, a pattern seen both in response to phonemes and tone-pips. Given the previous literature, it seems reasonable to posit that the greatly reduced P2 amplitude in RTT likely indexes severely affected automatic integration and retrieval of auditory stimulus statistics, and that this would diminish or even preclude effective processing of the auditory scene. The amplitude of this most severely affected component was below the minimal TD value in 9 out of 12 RTT subjects (75%), pointing to the clinical relevance of the P2 effect, and this clinical relevance is further bolstered by the significant correlation between P2 amplitude and the severity of Rett symptomatology, here somewhat crudely measured by the RSSS. The link between RSSS and auditory P2 might be explained by general neural abnormalities manifest in parallel in RTT. However they might be also be linked to some specific abnormalities in particular brain regions contributing to both effects. While RSSS is an integrative measure covering multiple domains, most of these can be linked to subcortical deficits, e.g. hypotonia, breathing irregularities, sleeping problems. Taking into account that the reticular-thalamo-cortical activating system was reported to contribute to the amplitude of the P2 (Velasco et al., 1989), both the clinical symptoms and P2 reduction in patients with RTT might be related to brainstem deficiencies.

It is worth pointing out that P2 abnormalities have also been reported in other clinical populations, such as atypical modulations of P2 by inter-stimulus interval (Jaffe-Dax et al., 2017) and stimulus statistics (Jaffe-Dax et al., 2015) in children with a diagnosis of dyslexia. However, the substantial P2 reduction seen here during passive listening seems to be specific to patients with RTT.

#### N2 attenuation (245-275 ms)

The next N2 component is generated in the vicinity of auditory cortex (Ponton et al., 2002) with potential additional sources in frontal cortex (Karch et al., 2010). This component has been related to the inhibition of irrelevant information, as it shows attenuation with stimulus repetition (Karhu et al., 1997) in children and young adults, and it is also reduced in elderly subjects compared to younger adults (Bertoli and Probst, 2005; Harris et al., 2012; Stothart and Kazanina, 2016). N2 reduction has also been reported in children with developmental dysphasia (Korpilahti and Lang, 1994) and other language impairments (Neville et al., 1993). Further supporting a relationship between N2 and language processing abnormalities, a longitudinal study found that decreased N2 amplitudes to non-speech stimuli between the ages of 4 and 8 years was associated with subsequent poorer word reading at school (Espy et al., 2004). As such, the current finding of robust N2 attenuation in patients with RTT may index deficient inhibitory processes that are a prerequisite for typical language development.

### Bridging animal and patient studies in RTT

Since the discovery that mutations of *MECP2* were causal to Rett Syndrome, this disease has been actively studied in pre-clinical animal models with targeted genetic manipulations (Guy et al., 2001; Shahbazian et al., 2002; Panayotis et al., 2011; Goffin et al., 2012). In addition to behavioral manifestations of Rett Syndrome, such as pronounced stereotypic forelimb motions, uncoordinated gait, reduced spontaneous movement, and irregular breathing, AEP components were also found to be abnormal in the rodent model of RTT (Goffin et al., 2012; Liao et al., 2012; Engineer et al., 2015). Our study bridges the animal and human research by examining similar AEP components to those studied in rodent models of RTT. The most pronounced effect of P2 amplitude reduction observed in our patients was also evident in RTT animal models: P2 amplitude was reduced both in rat and mice RTT models both in response to click and speech sounds (Goffin et al., 2012; Liao et al., 2012; Engineer et al., 2015). Consistent with the idea of the relevance of P2 to Rett symptomatology, P2 amplitude was typical in *Mecp2*-deficient mice before the first manifestation of symptoms (Goffin et al., 2012; Liao et al., 2012). Also in line with our results, N2 amplitude was significantly reduced in response to speech sounds in the rat model of RTT, the only model and condition where N2 was examined (Engineer et al., 2015). Noteworthy, in spite of evident attenuation of P2 and N2 components in *Mecp2*-deficient rats (Engineer et al., 2015), these rats learned to discriminate speech sounds similarly to wild-type animals. The mutant animals’ speech discrimination performance became significantly poorer than that of wild-types only in a more challenging condition that included background noise. Thus, the observed AEP abnormalities might not be crucial for simple auditory discrimination tasks but necessary for complex sound discrimination under noisy environmental conditions.

### Study Limitations

A limitation here was the broad age-range of participants, given that the AEP continues to mature across development over the age-range tested (Brandwein et al., 2011). Another issue was that we did not acquire hearing tests on the day of experimentation from RTT patients due to difficulties in assessing it in this population. However, the presence of prominent P1 and N1 components in the RTT group, clearly indicates that they could detect and decode auditory information.

### Summary

These data provide the first in-depth analysis of the early AEP in response to speech stimuli, revealing the absence of typical modulation of the P1 component by stimulus complexity in patients with Rett Syndrome. While the next N1 component was preserved in RTT, the later AEP components, P2 and N2, were almost completely abolished. The most severely affected component, the P2, was below TD levels in most RTT participants and was associated with symptomatology. Moreover, unlike other components, P2 was stable across development. As such, the P2 AEP component holds real promise as a neuromarker of RTT. It is especially noteworthy that P2 attenuation has also been reported in animal models of RTT, providing a potentially key translational link between this neuromarker in patients and RTT animal model studies.

## Acknowledgments

This work was supported by grants from the U.S. National Institute on Deafness and Other Communication Disorders (NIDCD R21 DC012447 to JJF and AD) and from The Rett Syndrome Research Trust (to JJF and SM). Work by OS was supported by a Fulbright scholarship. The authors thank the volunteers and their families, who all gave their time with great patience and grace to allow for the successful completion of this study. The authors acknowledge the participation of Mr. Douwe Horsthuis and Mr. Emmett Foxe during the data collection phase of this project.

